# Integrative physiological plasticity of *Agastache rugosa* (Fisch. & C.A.Mey.) Kuntze reveals complex adaptation to light and nutrient gradients

**DOI:** 10.1101/2024.10.01.616001

**Authors:** Khairul Azree Rosli, Azizah Misran, Latifah Saiful Yazan, Puteri Edaroyati Megat Wahab

## Abstract

We investigated the physiological plasticity of *Agastache rugosa* (Fisch. & C.A.Mey.) Kuntze, in response to different light and nutrient levels, demonstrating complex adaptive strategies. Through comprehensive analysis of leaf traits, photosynthetic parameters, and resource use efficiencies, we uncovered unexpected patterns in Rubisco dynamics and nutrient utilization, particularly in low-light conditions. *A. rugosa* exhibited subtle thermal regulation, maintaining relatively stable leaf temperatures across treatments through coordinated adjustments in morphology and gas exchange. Multivariate analyses showed distinct clustering of traits, underlining the integrated nature of plant responses to environmental heterogeneity. Leaf area increased under low-light conditions, while leaf mass area was higher in high-light environments, consistent with shade avoidance syndrome and leaf economics spectrum theory. Surprisingly, Rubisco concentration and use efficiency were generally increased under low light, suggesting a compensatory mechanism. Water use efficiency was higher in high-light conditions, but photosynthetic phosphorus and potassium use efficiencies displayed surprising increases under low light. The species exhibited tight coordination between CO_2_ supply and demand, as evidenced by strong correlations between net photosynthesis, stomatal conductance, and carboxylation efficiency. Our findings suggest that *A. rugosa* employs a suite of physiological and morphological adjustments to optimize resource acquisition and utilization that contribute to its ecological success.

**Highlight:** This study reveals complex adaptive strategies in *A. rugosa* under varying light and nutrient levels, showing unexpected patterns in Rubisco dynamics and nutrient use efficiencies while maintaining subtle thermal regulation across environments.

## Introduction

Plants exhibit remarkable plasticity in their ability to adapt to diverse environmental conditions, a trait that is crucial for their survival and productivity in various habitats. This adaptive capacity is particularly evident in their responses to light and nutrient availability, two of the most important factors influencing plant growth and development (Okunlola and Adelusi, 2014; Valladares *et al*., 2016; Sakuraba and Yanagisawa, 2018). Light is the main energy source for photosynthesis, while nutrients are essential building blocks for cellular structures and many metabolic processes. The interaction between light and nutrients shapes plant morphology, physiology, and biochemistry at multiple scales, from leaf-level traits to whole-plant resource allocation patterns (Freschet *et al*., 2018; Poorter *et al*., 2019). Understanding this complex interplay is fundamental to plant ecology and evolution and has critical implications for agriculture, forestry, and ecosystem management in the face of global environmental change (Elser *et al*., 2007; Grime and Pierce, 2012; Courbier and Pierik, 2019).

Extensive research has been conducted on how plants respond to light and nutrients, uncovering a range of adaptive mechanisms. Under low-light conditions, plants often exhibit shade avoidance or shade tolerance syndromes, characterized by increased leaf area, reduced leaf mass area (LMA), and altered pigment contents to maximize light capture (Lusk and Warton, 2007; Cagnola *et al*., 2012; Chitwood *et al*., 2015). Conversely, high-light environments often induce increased LMA, higher photosynthetic rates, and enhanced photoprotection (Coble and Cavaleri, 2014; Murchie and Ruban, 2020). Nutrient availability further modulates these light-mediated responses, with nutrient-rich conditions generally promoting greater investment in photosynthetic machinery and overall growth, while nutrient limitation often leads to conservative resource-use strategies (Aerts and Chapin III, 1999; Reich, 2014). However, recent studies have shown that plant responses to combinative light and nutrient gradients are not always predictable from their responses to each factor in isolation, highlighting the need for integrated approaches to understand plant adaptation strategies (Freschet *et al*., 2018; Bouain *et al*., 2019).

The physiological and biochemical mechanisms underlying plant responses to light and nutrients are intricate and multifaceted. At the leaf level, adaptation involves adjustments in photosynthetic capacity, often mediated through changes in Rubisco concentration and activation state, as well as modifications in light-harvesting complexes and electron transport (Hikosaka and Shigeno, 2009; Carmo-Silva *et al*., 2015). These changes are tightly connected with leaf morphology and anatomy alterations to optimize resource acquisition and utilization. Moreover, plants must balance carbon fixation with water loss through precise regulation of stomatal conductance, leading to trade-offs between water-use efficiency and photosynthesis (Flexas *et al*., 2016). Nutrient availability further complicates this process by affecting the allocation of resources to other physiological responses and structures. For instance, nitrogen and phosphorus play key roles in photosynthetic machinery and energy transfer, respectively, influencing photosynthetic nutrient use efficiencies (Hidaka and Kitayama, 2009; Yuan and Chen, 2015; You *et al*., 2018). Recent advances in our understanding of these processes have highlighted the importance of considering multiple resource limitations simultaneously (Poorter *et al*., 2012, page 20; Lambers *et al*., 2019).

Despite the wealth of information on plant responses to light and nutrients, important gaps remain in our grasp of how plants integrate multiple environmental signals to optimize their growth across heterogeneous landscapes. In particular, the mechanisms by which plants maintain physiological homeostasis, especially thermal regulation, under different levels of resource availability are not fully elucidated. Additionally, the plasticity of resource-use efficiencies across light and nutrient gradients and its implications for agricultural practices and ecosystem functioning require further investigation. To address these gaps, we conducted a comprehensive study on *Agastache rugosa* (Fisch. & C.A.Mey.) Kuntze, a versatile herb with medicinal and culinary applications (Hou *et al*., 2022). *A. rugosa* is a perennial native to temperate and subtropical climates. Cultivating the herb in the tropics poses challenges due to strong light and low native soil fertility (Amissah et al. 2024; Terán et al. 2024). Hence, we hypothesized that *A. rugosa* would exhibit coordinated plasticity in leaf traits, photosynthetic processes, and resource-use efficiencies under varying light and nutrient levels, with specific adaptations to maintain physiological performance in different environments. Our objective was to elucidate the physiological and morphological responses of *A. rugosa* to light and nutrient treatments, focusing on photosynthetic adaptation and the relationships among major traits related to resource use that contribute to its ecological plasticity. By integrating physiological measurements with multivariate statistical approaches, this study aims to add new insights into the adaptive strategies of plants in heterogenous environments and enhance our understanding of plant functional ecology.

## Materials and methods

### Plant materials

*A. rugosa* seeds were bought from a local supplier (WHT Wellgrow Seeds, Malaysia). The seeds were surface-sterilized for 1 min with 70% ethanol, followed by 5% sodium hypochlorite solution for 10 min, then rinsed thrice with distilled water. The seeds were sown in 200-cell trays containing a 1:1 mixture of blonde peat (Pindstrup Mosebrug A/S, Denmark) and perlite. The seed cell trays were placed on capillary mats under white fluorescent light/dark (16/8 h) photoperiod (Park *et al*., 2020) in a room maintained at 20–25 °C and 50–70% relative humidity. The substrate was kept moist but not drenched. Seeds were allowed to germinate and grow under these conditions.

### Experimental setup

This experiment was conducted under two north-south oriented polytunnels (10 m long x 6 m wide x 2 m high) with both end-walls and side-walls (1.5 m high) at Farm 10, Faculty of Agriculture, Universiti Putra Malaysia. The polytunnels were enclosed with a single layer of clear polyethylene film (280 microns thick). The end-walls were open, and side-walls were rolled down during the experimental period. To simulate low-light conditions, a tunnel roof was covered externally with a single layer of 50% black netting (Supplementary Fig. S1). Photosynthetic photon flux density (PPFD), air temperature (T_Air_), and relative humidity (RH) data were recorded using a light meter (LI-189, LI-COR, Lincoln, NE, USA) and dataloggers (Supplementary Fig. S2). Six-week-old seedlings of uniform size were transplanted to black polythene pots (15 cm diameter x 25 cm high) containing ≈3.5 kg pot^-1^ (≈24 cm depth) of commercial 3:2:1 potting mix. Details on the chemical properties of the potting mix are listed in Table S1. Four nutrient levels based on NPK (16:16:16); low (NPK1, 40 mg kg^-1^, 1.26 g), moderate (NPK2, 80 mg kg^-1^, 2.52 g), high (NPK3, 120 mg kg^-1^, 3.78 g) and very high (NPK4, 160 mg kg^-1^, 5.04 g) were nested under two light levels; high- light (HL, 0% shade) and low-light (LL, 50% shade), and arranged in randomized complete block design with four replicates in two sets (4 replicates x 8 treatment combinations x 2 sets = 64 pots).

Light treatments were initiated at transplanting or 0 days after transplanting, while nutrients were split-applied 6, 13, 27, 41, 55, and 69 days after transplanting. Compound fertilizer (YaraMila, Norway) was used as source of nutrients, manually applied in individual holes (5 cm deep). Plants were watered after each nutrient application via drip irrigation.

**Supplementary Fig. S1.**
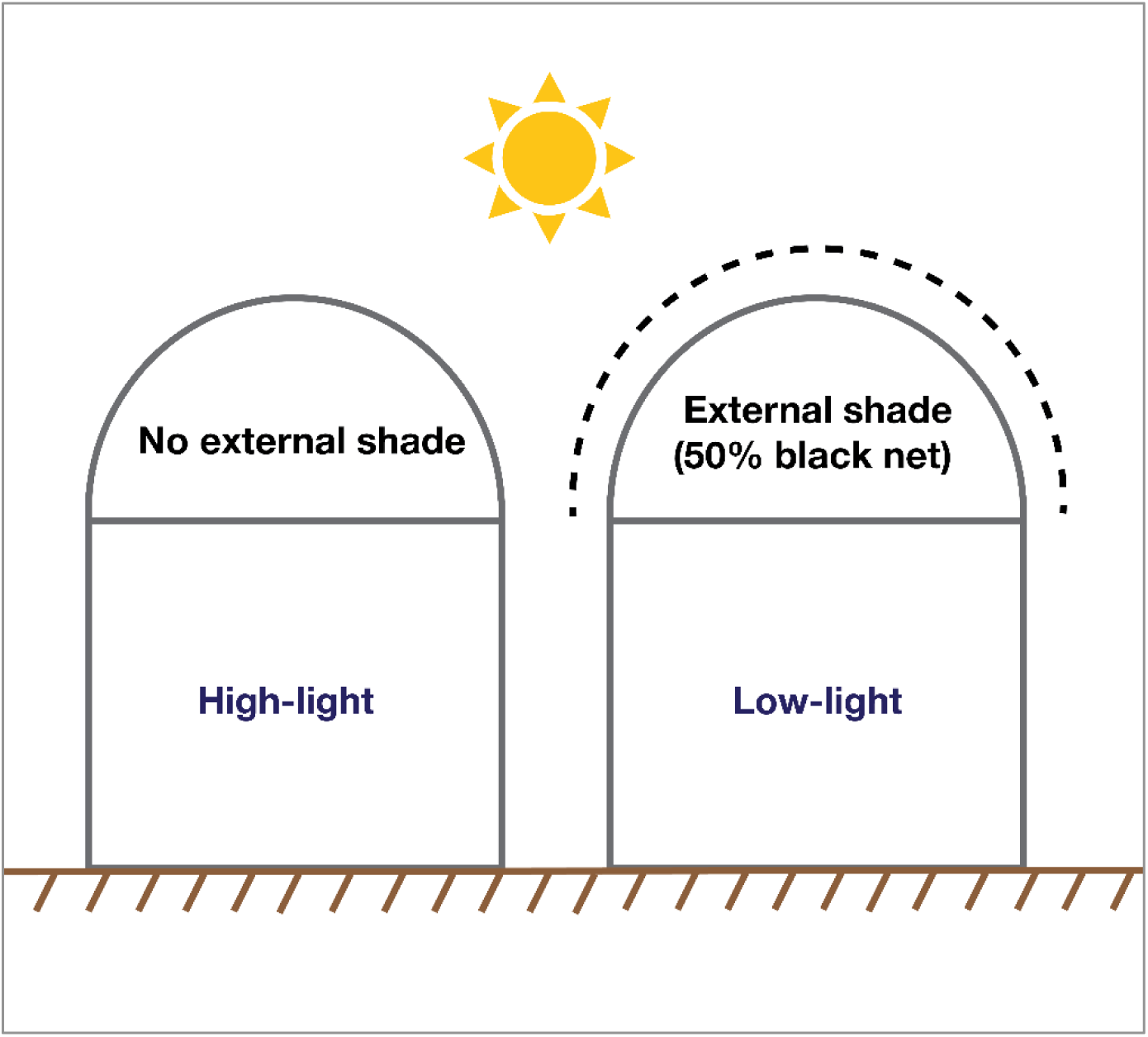
Setup of the polytunnels for the two light treatments.

**Supplementary Fig. S2.**
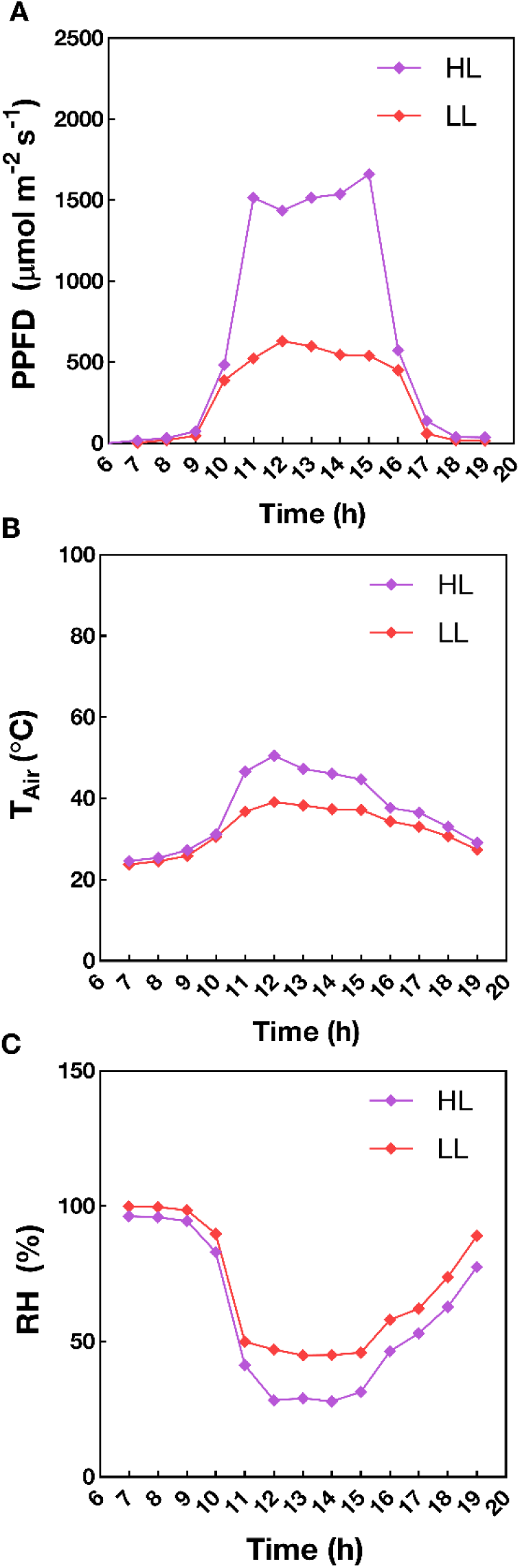
Diurnal variations of microclimatic parameters in tunnels set for high-light (HL, 0% shade) and low-light (LL, 50% shade) levels, averaged over September to December 2020. Abbreviations: PPFD, photosynthetic photon flux density; T_Air_, air temperature; RH, relative humidity.

**Table S1.** Chemical properties of potting mix.

| Properties | Value |
| --- | --- |
| pH (H<sub>2</sub>O) | 6.90 |
| Cation exchange capacity | 11.80 |
| Organic carbon (C) (%) | 6.50 |
| Total nitrogen (N) (%) | 0.14 |
| C/N ratio | 47.00 |
| Total phosphorus (P) (mg kg<sup>-1</sup>) | 612.12 |
| Available phosphorus (P) (mg kg<sup>-1</sup>) | 551.00 |
| Exchangeable cations (meq100 g<sup>-1</sup>) |  |
| Sodium (Na) | 0.35 |
| Potassium (K) | 2.97 |
| Calcium (Ca) | 3.34 |
| Magnesium (Mg) | 0.82 |
| Available microelements (mg kg<sup>-1</sup>) |  |
| Boron (B) | 1.16 |
| Iron (Fe) | 43.39 |
| Copper (Cu) | 0.68 |
| Zinc (Zn) | 4.25 |
| Manganese (Mn) | 27.59 |

### Gas exchange and water use efficiency

Gas exchange measurements were taken 84 days after transplanting between 0900 h and 1030 h. Light-saturated net photosynthesis rates (A, µmol CO_2_ m^−2^ s^−1^), transpiration rate (E, mmol H_2_O m^2^ s^−1^), stomatal conductance (g_sw_, mol H_2_O m^2^ s^−1^), intercellular CO_2_ (Ci, μmol mol^-1^), and leaf temperature (T_Leaf_, °C) were measured on the third or fourth youngest fully expanded leaf from the shoot apex of using a photosynthesis instrument (LI-6800, LI-COR Incorporated, Lincoln, NE, USA) fitted with a 6 cm^2^ leaf chamber with controlled light and gas flow (6800-01A). The device was calibrated and set to measure at an ambient CO_2_ concentration of 400 μmol mol^-1^, 50–60% relative humidity, 1500 μmol m^-2^ s^-1^ PPFD, and airflow rate of 500 μmol air s^-1^. Data were recorded at steady-state conditions in between measurements. Carboxylation efficiency (CE, µmol m^−2^ s^−1^) was determined by dividing A with Ci (Rymbai *et al*., 2014). Water use efficiency (WUE) indexes were estimated based on Hatfield and Dold (2019). Instantaneous WUE (WUE_ins_, µmol CO_2_ mmol H_2_O^−1^) was calculated as A divided by E, whereas intrinsic WUE (WUE_int_, µmol CO_2_ mol H_2_O^−1^) was determined by dividing A with g_sw_.

### Leaf trait measurements

After gas exchange measurements, leaves were harvested and brought to the laboratory. Leaf area (LA, cm^2^) of detached leaves was measured using a leaf area meter (LI-3100C, LI-COR, Lincoln, NE, USA), and their fresh weights (g) were recorded. Leaf dry weights (g) were measured after drying at 105 °C for 30 min, followed by 80°C for two days in a forced-air oven (Memmert UNB- 500, Germany). Leaf mass area was calculated, which is the amount of leaf dry mass per unit leaf area (LMA, mg cm^-2^), and leaf water content, the amount of water per unit leaf dry weight (LWC, %) (Adet *et al*., 2024; Guo *et al*., 2024).

### Nutrient analysis

Dried fully expanded leaves were powdered in an electric stainless-steel grinder (Nima NM-8300, Japan) to pass a 300-mesh sieve (Motsara and Roy, 2008). Briefly, 0.25 g of ground samples were liquefied through wet acid digestion with 5 mL of sulfuric acid and hydrogen peroxide. The digest solutions were diluted to 1:10 (v/v) and tested for total nitrogen calorimetrically (Koistinen *et al*., 2019), while phosphorus and potassium levels were analyzed using atomic absorption spectrometry (model 5100, PerkinElmer, USA). Photosynthetic nitrogen, phosphorus, and potassium use efficiencies (PNUE, µmol CO_2_ mol N^−1^ s^−1^; PPUE, µmol CO_2_ mol P^−1^ s^−1^; PKUE, µmol CO_2_ mol K^−1^ s^−1^) were calculated according to Prieto et al. (2023) as follows:

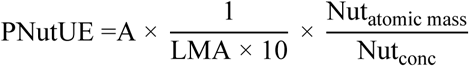

Where A is the net photosynthesis rate (µmol CO_2_ m^−2^ s^−1^), LMA is the leaf mass per area (mg cm^−2^), Nut_content_ is the leaf nutrient concentration (mg g^−1^), and Nut_atomic mass_ is the atomic mass of nutrient (14.61 g mol^−1^ for N, 30.974 g mol^−1^ for P and 39.1 g mol^−1^ for K).

### Rubisco

Rubisco (EC 4.1.1.39) concentration was measured via sodium dodecyl sulfate-polyacrylamide gel electrophoresis (SDS–PAGE) method with some modifications (Xiong *et al*., 2017). The third or fourth youngest fully expanded leaves from the apex were flash-frozen at harvest and powdered with liquid nitrogen using precooled porcelain mortars and pestles. Frozen samples were weighed into 2-mL microtubes at 0.1 g each and homogenized with 1 mL of cold extraction buffer (5 mM of 2-mercaptoethanol and 12.5% (v/v) glycerol in 50 mM Tris-HCL buffer pH 8.0). Samples were centrifuged at 4 °C for 10 min at 8,000x*g.* Supernatants were added with equal volume of loading buffer (2% SDS, 4% 2-mercaptoethanol, and 10% glycerol in distilled water) into fresh microtubes and boiled in a water bath for 5 min. SDS-PAGE was performed using a Mini PROTEAN® Tetra Cell system (BioRad, Hercules, CA, USA). Samples and pre-stained protein standard (SeeBlue™ Plus 2, Invitrogen, Carlsbad, CA, USA) were loaded into the wells at 10 µL and 5 µL respectively, were loaded onto SDS-PAGE gel containing 4% (w/v) stacking gel, and 12.5% (w/v) separating gel. The gel electrophoresis was run at 100V for about 90 min. After electrophoresis, the gels were rinsed several times with deionized water and stained with CBB Stain One Super (Nacalai Tesque, Inc., Kyoto, Japan) overnight while shaken at 40 rpm. The gels were then destained with ultrapure water until the background was colorless. Both large (RbcL, ∼55 kDa) and small (RbcS, ∼14 kDa) subunits, and background gel blank were excised and transferred into microtubes and added with 2 mL of formamide. Samples and blank were incubated at 50°C in an oven for 2 h. Absorbance was read at 595 nm, and the values were compared against a standard curve prepared using BSA solutions. Rubisco concentration was expressed as milligrams per gram tissue fresh weight (mg/g FW). Rubisco use efficiency and Rubisco nitrogen fraction were estimated based on Hikosaka and

Shigeno (2009). Rubisco use efficiency, or the photosynthetic rate per unit Rubisco (RBUE, µmol CO_2_ g^−1^ s^−1^), was calculated as follows:

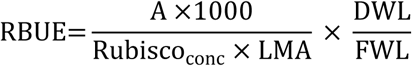

Where A is the net photosynthesis rate (µmol CO_2_ m^−2^ s^−1^), LMA is the leaf mass per area (g m^−2^), Rubisco_conc_ is the concentration of Rubisco per gram tissue fresh weight (mg g^-1^ FW), DWL is the leaf dry weight (g) and FWL is the leaf fresh weight (g). Rubisco nitrogen fraction (RNF) was calculated based as follows:

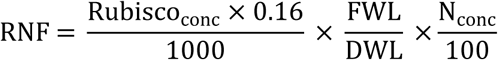

Where Rubisco_conc_ is the concentration of Rubisco per gram tissue fresh weight (mg g^-1^ FW), DWL is the leaf dry weight (g), FWL is the leaf fresh weight (g) and N_conc_ is the total leaf N concentration (%).

### Statistical analysis

Data were analyzed using SAS^®^ version 9.4 by the general linear model (PROC GLM). Wherever necessary, data was transformed before analysis through Box-Cox transformations to ensure the normality of residuals was satisfied. A combined analysis of variance was performed for the four nutrient levels nested under light levels (Bowley, 1999; Moore and Dixon, 2015). The means of significant main effects and interactions were separated with the least significant difference (LSD) posthoc test (Vargas *et al*., 2015). Pearson correlation test and principal component analysis (PCA) were used to explore the relationship among variables. Graphs, heatmap and biplot after PCA were made using OriginPro^®^ 2024b.

## Results

### Leaf traits

Leaf traits of *A. rugosa* exhibited significant variations (P < 0.05) under different light and nutrient levels (Fig. 1). Leaf area (LA) displayed a general increasing trend with higher nutrient levels for both high-light (HL) and low-light (LL) treatments (Fig. 1A). LA was consistently larger under LL compared to HL conditions across all nutrient levels, with the highest values observed in HL- NPK4 (1671.07 cm² plant⁻¹) and LL-NPK4 (1542.00 cm² plant⁻¹) treatments. Leaf mass per area (LMA) exhibited an opposite trend, with HL conditions resulting in higher LMA values across all nutrient levels (Fig. 1B). The highest LMA was observed in HK-NPK1 (6.32 mg cm⁻²), while LL conditions consistently demonstrated lower LMA values. Leaf water content (LWC) was generally higher under LL conditions compared to HL (Fig. 1C). The most pronounced difference in LWC between light treatments occurred in NPK4, with LL showing significantly higher LWC (80.13%) compared to HL (69.37%). LL-NPK1 exhibited the highest LWC (84.22%) among all treatments.

**Fig. 1.**
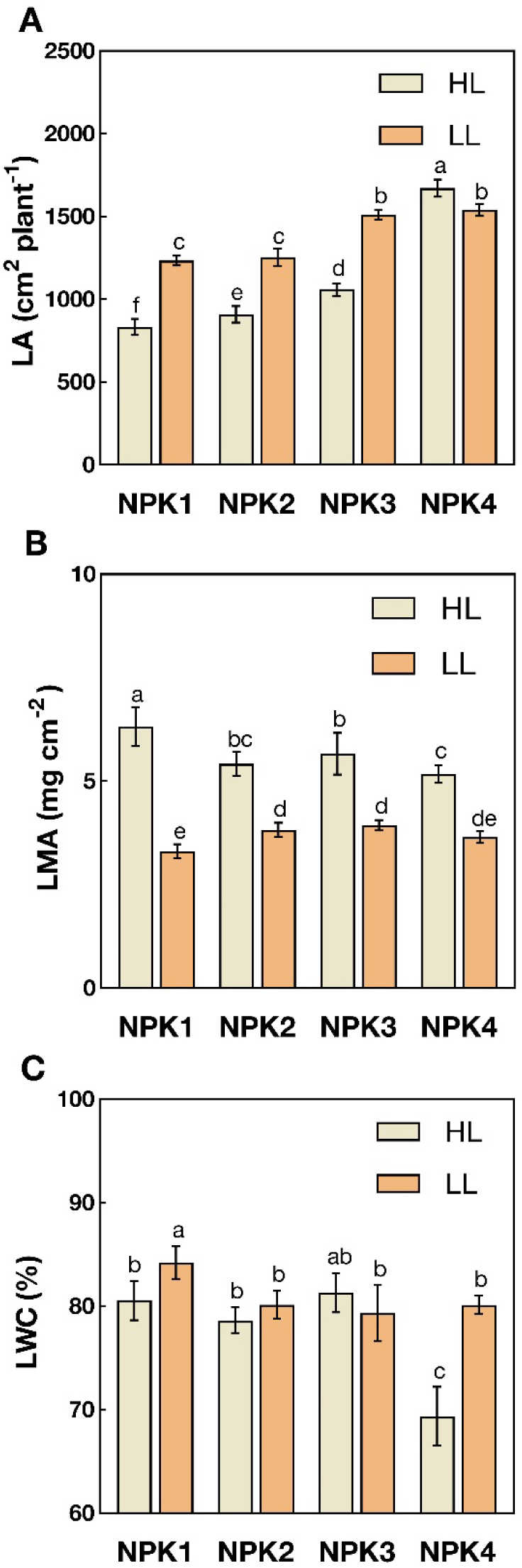
Leaf traits of *A. rugosa* under different light and nutrient levels. Abbreviations: HL, high-light (0% shade); LL, low-light (50% shade); NPK1, low-nutrient (40 mg kg^-1^); NPK2, moderate-nutrient (80 mg kg^-1^); NPK3, high-nutrient (120 mg kg^-1^); NPK4, very high-nutrient (160 mg kg^-1^); LA, leaf area; LMA, leaf mass area; LWC, leaf water content. The vertical bars are mean ± SD (n = 4). Different letter(s) above bars indicate significant differences according to LSD. LSD = Least significant difference; SD = Standard deviation. Each value corresponds to the mean of four biological replicates.

### Gas exchange and carboxylation efficiency

Interaction effects (P < 0.05) were observed between different light and nutrient levels on leaf gas exchange and carboxylation efficiency of *A. rugosa* (Fig. 2). Net photosynthetic rate (A) showed a marked increase under high-nutrient levels (NPK3 and NPK4), with the highest values observed in HL-NPK4 (17.62 µmol CO_2_ m^-2^ s^-1^) treatment followed by LL-NPK3 (Fig 2. A). Transpiration rate (E) showed a similar trend, with NPK4 treatments yielding the highest rates, particularly under LL-NPK4 (7.21 mmol H_2_O m^-2^ s^-1^) conditions (Fig 2B). Stomatal conductance (g_sw_) mirrored this pattern, peaking in LL-NPK4 (0.32 mol H_2_O m^-2^ s^-1^) treatment (Fig 2C). Interestingly, intercellular CO_2_ concentration (Ci) showed less variation across treatments, though a slightly increasing trend was found with higher nutrient levels (Fig 2D). Leaf temperature (T_Leaf_) showed minimal variation across treatments, with values ranging between approximately 34°C and 37°C. However, a slight increase in T_Leaf_ was observed in higher nutrient levels, especially under LL conditions (Fig 2E). Carboxylation efficiency (CE) showed a significant increase in NPK3 and NPK4 treatments, with the highest efficiency observed under HL-NPK3 (68.67 µmol m^-2^ s^-1^) conditions (Fig 2F).

**Fig. 2.**
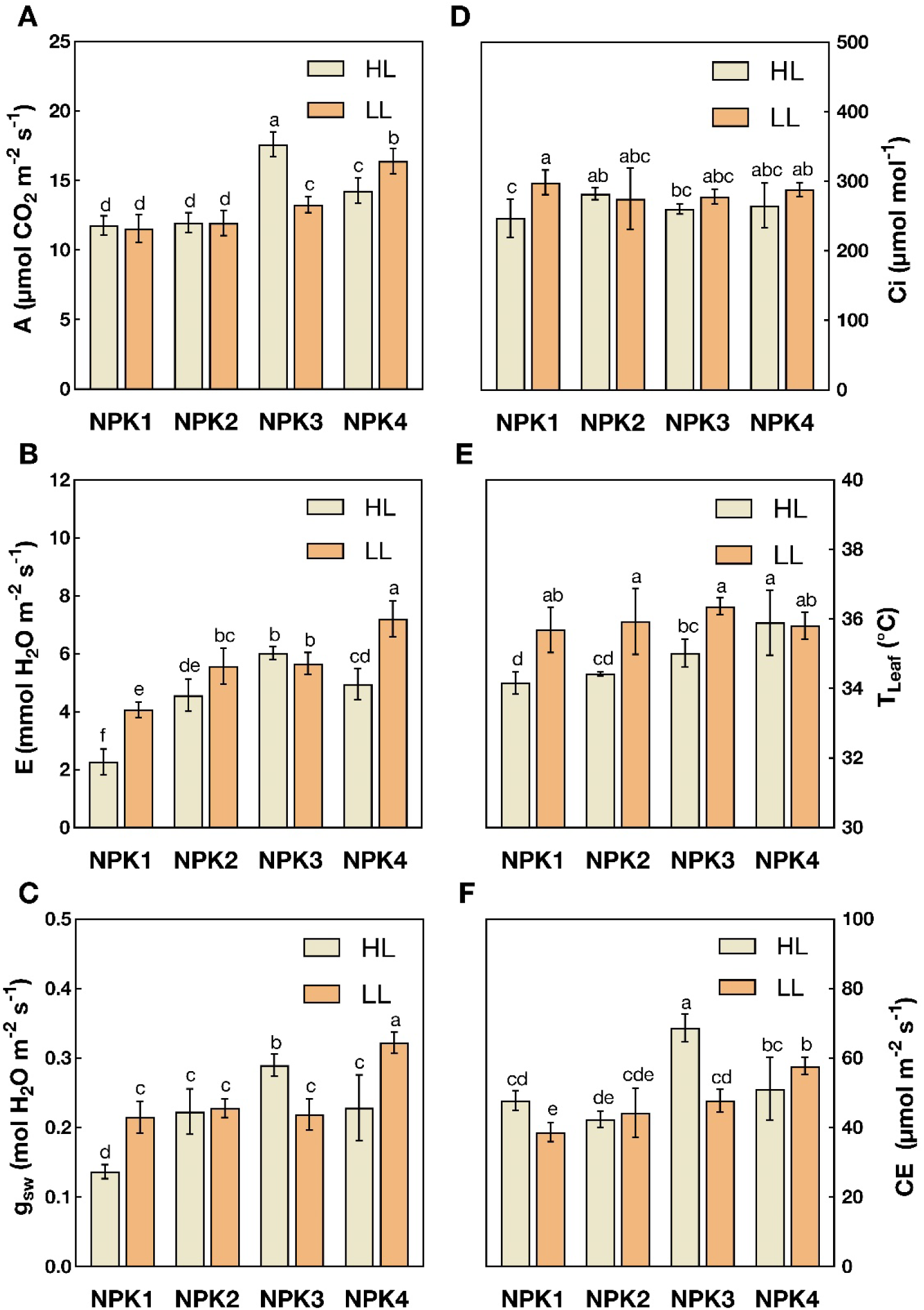
Leaf gas exchange and carboxylation efficiency of *A. rugosa* under different light and nutrient levels. Abbreviations: HL, high-light (0% shade); LL, low-light (50% shade); NPK, low-nutrient (40 mg kg^-1^); NPK2, moderate-nutrient (80 mg kg^-1^); NPK3, high-nutrient (120 mg kg^-1^); NPK4, very high- nutrient (160 mg kg^-1^); A, net photosynthetic rate; E, transpiration rate; g_sw_, stomatal conductance; Ci, intercellular CO_2_; T_Leaf_, leaf termperature; CE, carboxylation efficiency. The vertical bars are mean ± SD (n = 4). Different letter(s) above bars indicate significant differences according to LSD. LSD = Least significant difference; SD = Standard deviation. Each value corresponds to the mean of four biological replicates.

### Rubisco content and utilization

Light and nutrient significantly influenced (P < 0.05) Rubisco content and utilization in *A. rugosa* (Fig. 3). Rubisco concentration (Fig. 3A) generally increased with higher nutrient levels, with HL- NPK4 treatment showing the highest value (7.98 mg g^-1^ FW). Intriguingly, LL conditions resulted in higher Rubisco concentrations than HL conditions across all nutrient levels except for NPK3 and NPK4. Rubisco use efficiency (RBUE, Fig. 3B) displayed less pronounced differences across treatments, with the most notable increase observed in the LL-NPK4 condition (13.75 μmol CO_2_ g^-1^ s^-1^). Rubisco nitrogen fraction (RNF, Fig. 3C) demonstrated the most striking difference in the LL-NPK2 condition, reaching a peak of 1.19, markedly higher than all other treatments. Generally, LL conditions resulted in higher RNF values across all nutrient levels.

**Fig. 3.**
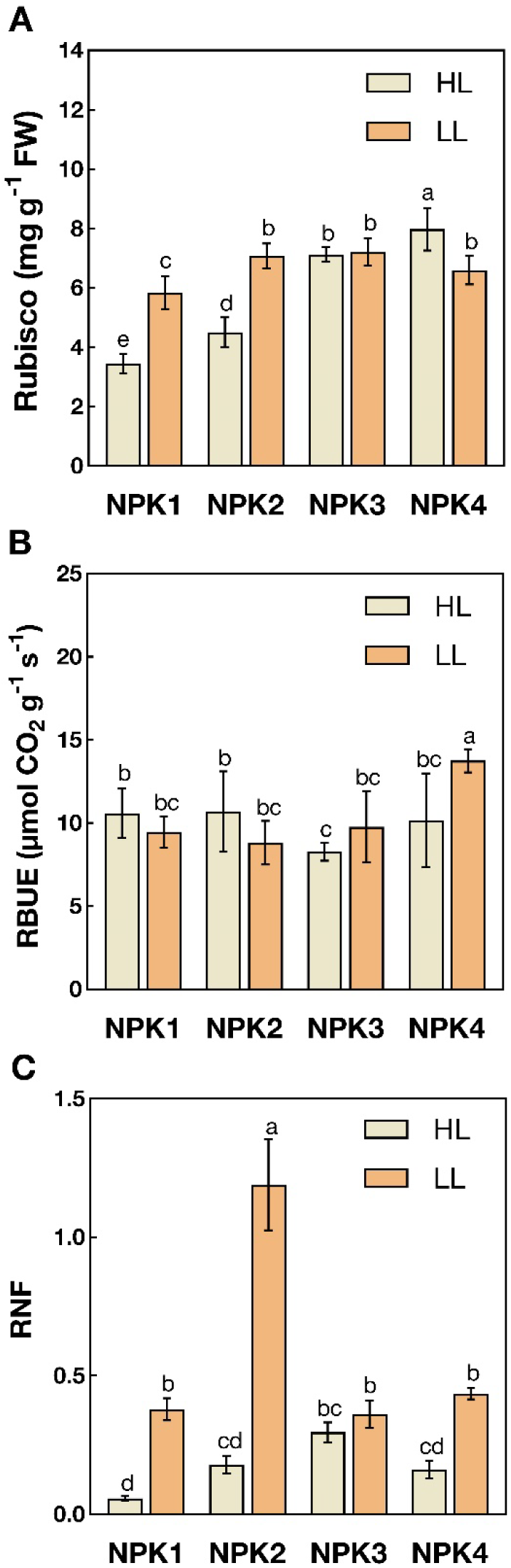
Rubisco content and utilization in *A. rugosa* under different light and nutrient levels. Abbreviations: HL, high-light (0% shade); LL, low-light (50% shade); NPK1, low-nutrient (40 mg kg^-1^); NPK2, moderate-nutrient (80 mg kg^-1^); NPK3, high-nutrient (120 mg kg^-1^); NPK4, very high-nutrient (160 mg kg^-1^); RBUE, Rubisco use efficiency; RNF, Rubisco nitrogen fraction. The vertical bars are mean ± SD (n = 4). Different letter(s) above bars indicate significant differences according to LSD. LSD = Least significant difference; SD = Standard deviation. Each value corresponds to the mean of four biological replicates.

### Water use efficiency

Light and nutrients significantly affected (P < 0.05) the water use efficiency (WUE) indexes of *A. rugosa* (Fig. 4). Plants grown under HL conditions exhibited markedly higher instantaneous WUE (WUE_ins_) than those under LL conditions (Fig. 4A). When examining the effects of nutrient levels (Fig. 4B), the low-nutrient treatment (NPK1) had the highest WUE_ins_ (4.10 µmol CO_2_ mmol^-1^ H_2_O, which was significantly different from all other nutrient treatments (NPK2, NPK3 and NPK4) that showed similar, lower WUE_ins_ values. Intrinsic WUE (WUE_int_) displayed a more complex pattern across treatments (Fig. 4C). Under NPK1, HL conditions led to markedly higher WUE_int_ compared to LL. However, this light-dependent difference was not observed in the other nutrient treatments. Interestingly, NPK2 and NPK3 treatments exhibited intermediate WUE_int_ values that did not differ significantly between HL and LL conditions.

**Fig. 4.**
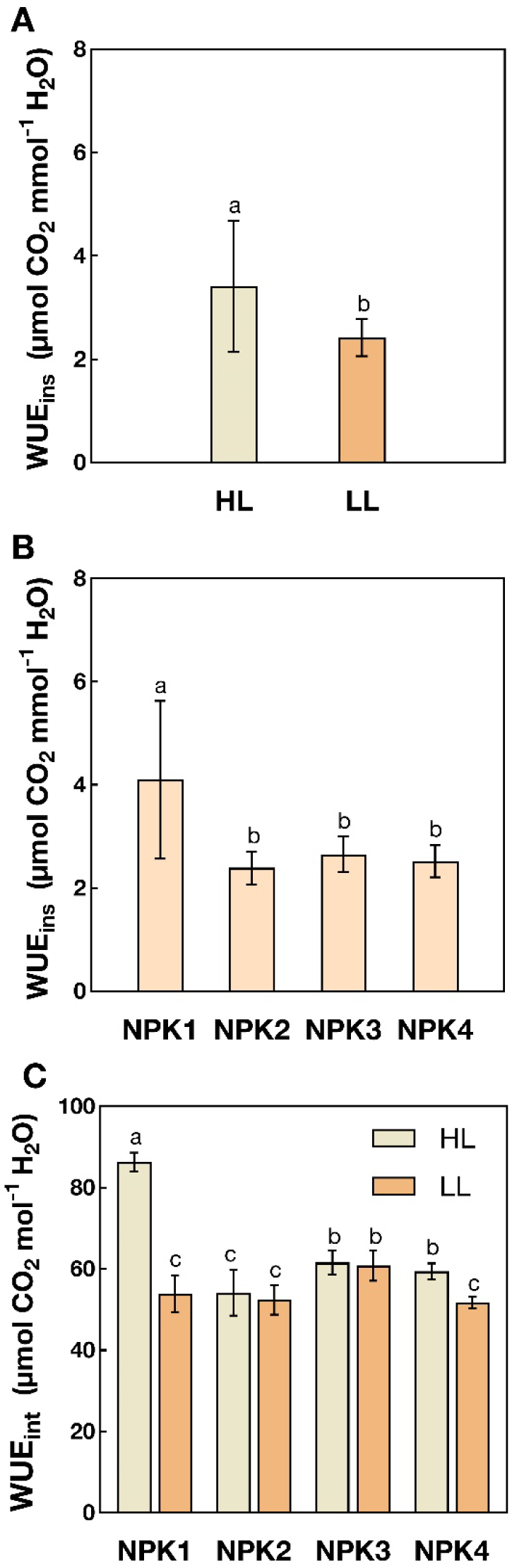
Changes in water use efficiency of *A. rugosa* under varying light and nutrient levels. Abbreviations: HL, high-light (0% shade); LL, low-light (50% shade); NPK1, low- nutrient (40 mg kg^-1^); NPK2, moderate-nutrient (80 mg kg^-1^); NPK3, high-nutrient (120 mg kg^-1^); NPK4, very high-nutrient (160 mg kg^-1^); WUE_ins_, instantaneous water use efficiency; WUE_int_, intrinsic water use efficiency. The vertical bars are mean ± SD (n = 4). Different letter(s) above bars indicate significant differences according to LSD. LSD = Least significant difference; SD = Standard deviation. Each value corresponds to the mean of four biological replicates.

### Photosynthetic nutrient use efficiency

Photosynthetic nutrient use efficiency of *A. rugosa* varied significantly (P < 0.05) under different light and nutrient levels. Photosynthetic nitrogen use efficiency (PNUE) was significantly higher under HL (98.64 µmol CO_2_ mol N^-1^ s^-1^) compared to LL (65.80 µmol CO_2_ mol N^-1^ s^-1^) conditions (Fig. 5A). Furthermore, PNUE showed a complex response to nutrient levels, with NPK2 having the lowest value (41.94 µmol CO_2_ mol N^-1^ s^-1^) while NPK4 had the highest (92.02 µmol CO_2_ mol N^-1^ s^-1^) efficiency (Fig. 5B). Photosynthetic phosphorus use efficiency (PPUE) generally increased with nutrient levels, with the highest values found in the LL-NPK4 (42.29 µmol CO_2_ mol P^-1^ s^-1^) treatment (Fig. 5C). Photosynthetic potassium use efficiency (PKUE) exhibited a similar pattern to PPUE, with LL conditions yielding higher efficiencies across all nutrient treatments (Fig. 5D). PKUE also showed an increasing trend with nutrient levels, reaching its maximum at LL-NPK4 (384.36 µmol CO_2_ mol K^-1^ s^-1^).

**Fig. 5.**
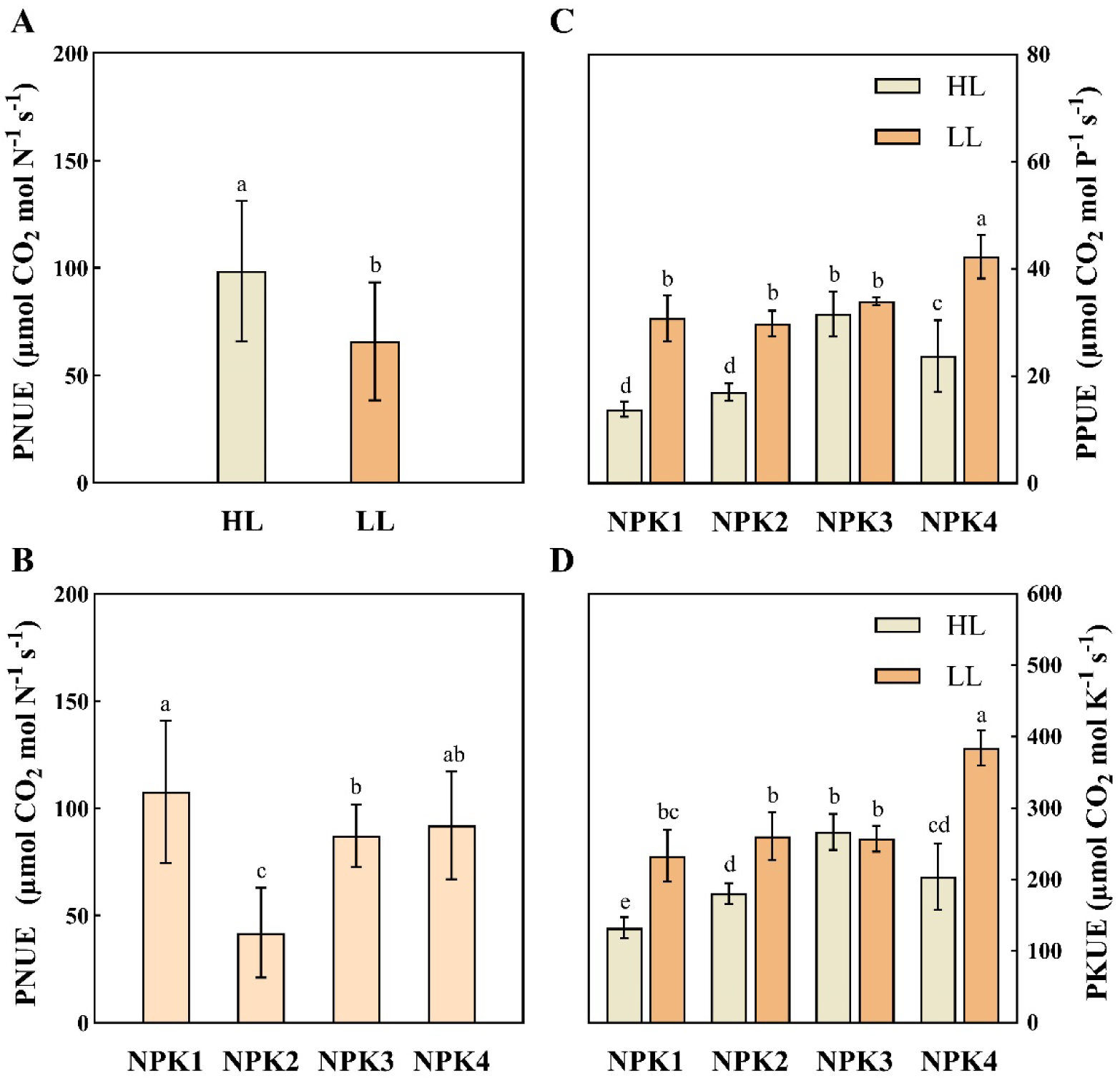
Changes in photosynthetic nutrient use efficiency of *A. rugosa* under different light and nutrient levels. Abbreviations: HL, high-light (0% shade); LL, low-light (50% shade); NPK1, low-nutrient (40 mg kg^-1^); NPK2, moderate-nutrient (80 mg kg^-1^); NPK3, high-nutrient (120 mg kg^-1^); NPK4, very high-nutrient (160 mg kg^-1^); PNUE, photosynthetic nitrogen use efficiency; PPUE, photosynthetic phosphorus use efficency. PKUE, photosynthetic potassium use efficiency. The vertical bars are mean ± SD (n = 4). Different letter(s) above bars indicate significant differences according to LSD. LSD = Least significant difference; SD = Standard deviation. Each value corresponds to the mean of four biological replicates.

**Fig. 6.**
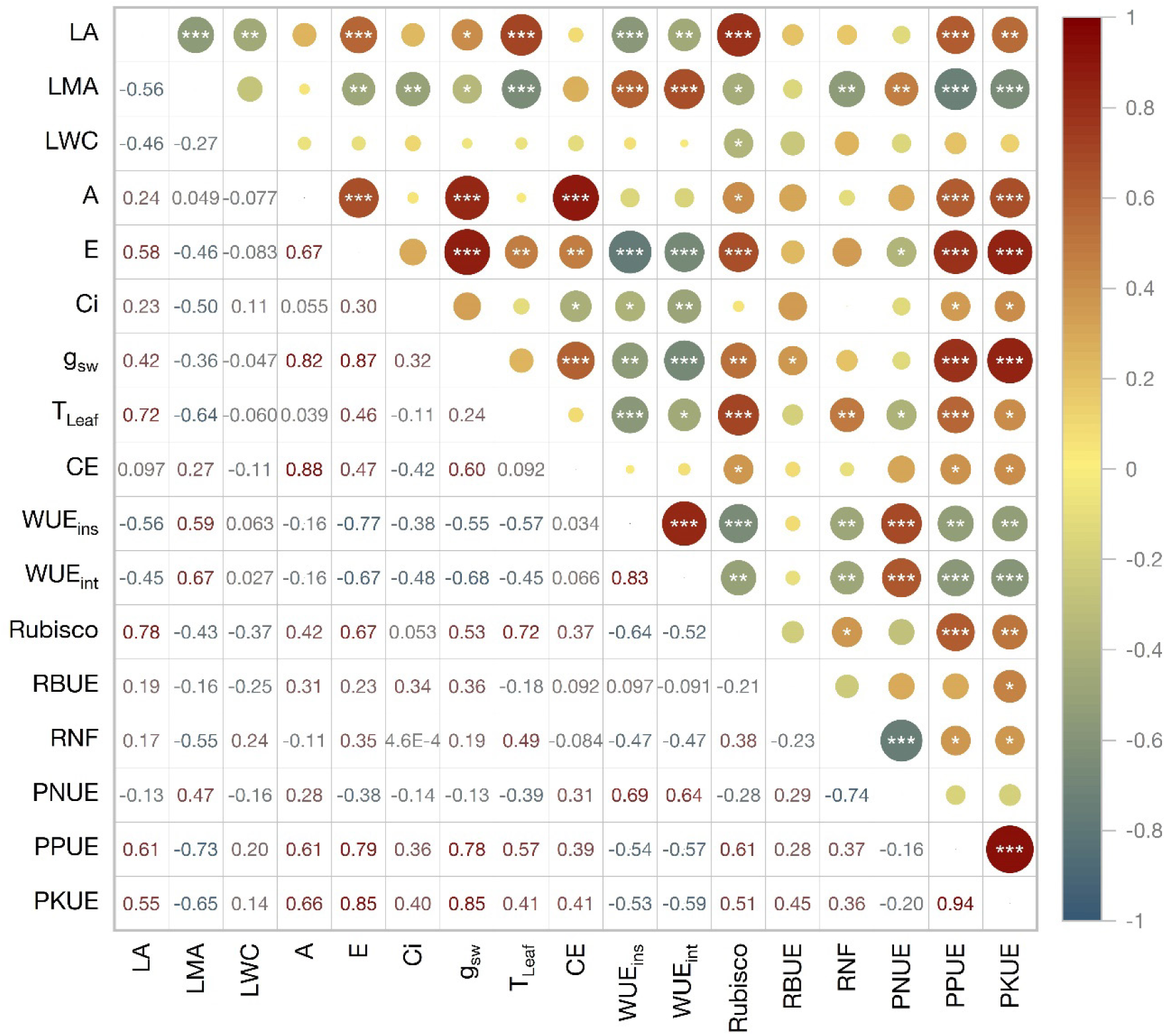
Heatmap showing correlations among traits related to physiological plasticity and resource use in *A. rugosa* under different light and nutrient levels. Abbreviations: LA, leaf area; LMA, leaf mass area; LWC, leaf water content; A, net photosynthesis rate; E, transpiration rate; Ci, intercellular CO_2_; g_sw_, stomatal conductance; T_Leaf_, leaf temperature; CE, carboxylation efficiency; WUE_ins_, instantaneous water use efficiency; WUE_int_, intrinsic water use efficiency; RBUE, Rubisco use efficiency; RNF, Rubisco nitrogen fraction; PNUE, photosynthetic nitrogen use efficiency; PPUE, photosynthetic phosphorus efficiency; PKUE, photosynthetic potassium use efficiency (∗∗∗P < 0.001; ∗∗P < 0.01; ∗P < 0.05).

### Correlations among measured traits

Pearson correlation analysis reveals complex relationships among traits related to resource use in *A. rugosa* under different light and nutrient levels. Leaf area (LA) had strong positive correlations with transpiration rate (E) (r = 0.58, P < 0.001), leaf temperature (T_Leaf_) (r = 0.72, P < 0.001), and Rubisco concentration (r = 0.78, P < 0.001). Leaf mass area (LMA) was negatively correlated with several traits, including LA (r = -0.56, P < 0.001) and transpiration rate (E) (r = -0.46, P < 0.01). Net photosynthetic rate (A) had robust positive associations with stomatal conductance (g_sw_) (r = 0.82, P < 0.001) and carboxylation efficiency (CE) (r = 0.88, P < 0.001). Notably, photosynthetic phosphorus use efficiency (PPUE) and photosynthetic potassium use efficiency (PKUE) were both highly correlated (r = 0.94, P < 0.001), showing strong positive relationships with E, g_sw,_ and A. Instantaneous water use efficiency (WUE_ins_) and intrinsic water use water use efficiency (WUE_int_) displayed a strong positive correlation (r = 0.83, P < 0.001) while being negatively associated with most other parameters, particularly E and g_sw_. Rubisco use efficiency (RBUE) exhibited weak to moderate correlations with most traits, with the strongest positive correlation observed with Ci (r = 0.34, P < 0.05). Rubisco nitrogen fraction (RNF) showed a significant positive correlation with T_Leaf_ (r = 0.49, P < 0.001) and negative correlations with LMA (r = -0.55, P < 0.001) and water use efficiency indexes.

### Principal component analysis of measured traits

The PCA biplot reveals distinct clustering patterns in traits related to resource use in *A. rugosa* under different light and nutrient levels (Fig. 7). The first two principal components accounted for 62.9% of the total variation (Dim1: 43.8%, Dim2: 19.1%). High-light treatments (HL-NPK1 to HL-NPK4) were predominantly associated with higher values for leaf mass area (LMA), water use efficiency indexes (WUE_ins_ and WUE_int_), and photosynthetic nitrogen use efficiency (PNUE). In contrast, low-light treatments (LL-NPK1 to LL-NPK4) correlated more strongly with increased leaf area (LA), net photosynthesis (A), carboxylation efficiency (CE), and Rubisco use efficiency (RBUE). Nutrient levels also affected trait distribution, with higher nutrient levels associated with higher photosynthetic rates and resource use efficiencies. Interestingly, leaf water content (LWC) and Rubisco nitrogen fraction (RNF) showed opposing relationships, with LWC aligning more closely with high-light treatments and RNF with low-light treatments. The biplot further illustrated complex interactions between physiological traits, with clusters of closely related parameters such as photosynthetic potassium use efficiency (PKUE), transpiration rate (E), and photosynthetic phosphorus use efficiency (PPUE).

**Fig. 7.**
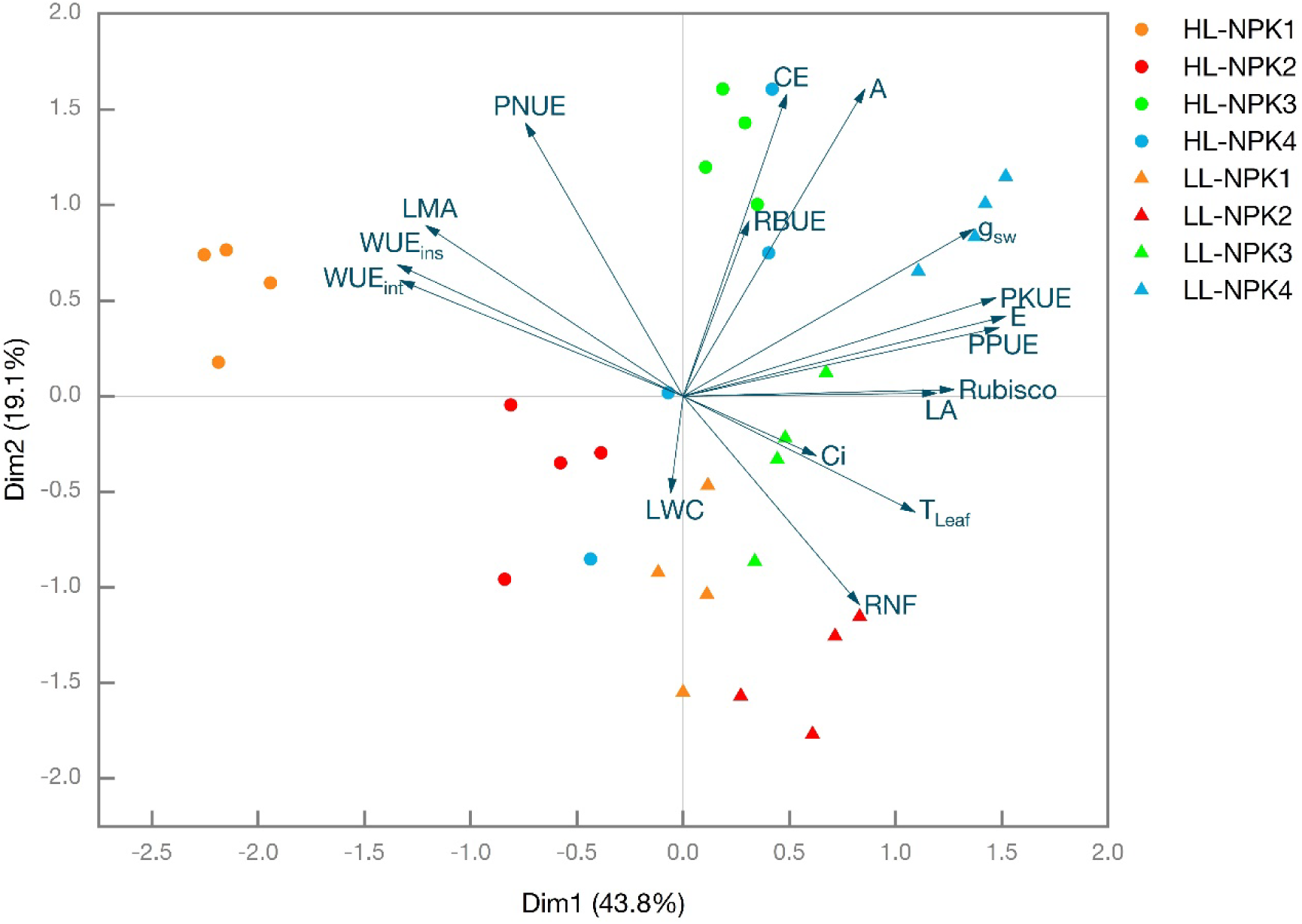
Principal component analysis (PCA) biplot displaying traits related to physiological plasticity and resource use in *A. rugosa* under varying light and nutrient levels. Abbreviations: HL, high-light (0% shade); LL, low-light (50% shade); NPK, low-nutrient (40 mg kg^-1^); NPK2, moderate-nutrient (80 mg kg^-1^); NPK3, high-nutrient (120 mg kg^-1^); NPK4, very high-nutrient (160 mg kg^-1^); LA, leaf area; SLW, specific leaf weight; LWC, leaf water content; A, net photosynthesis rate; E, transpiration rate; Ci, intercellular CO_2_; g_sw_, stomatal conductance; T_Leaf_, leaf temperature; CE, carboxylation efficiency; WUE_ins_, instantaneous water use efficiency; WUE_int_, intrinsic water use efficiency; RBUE, Rubisco use efficiency; RNF, Rubisco nitrogen fraction; PNUE, photosynthetic nitrogen use efficiency; PPUE, photosynthetic phosphorus efficiency; PKUE, photosynthetic potassium use efficiency.

### Cluster analysis of selected traits

The HCA dendrogram demonstrates three distinct clusters of variables that largely correspond to the spatial groupings observed in the PCA biplot (Fig. 8). The largest cluster, highlighted in pink, comprises a diverse set of parameters including leaf area (LA), Rubisco content, leaf temperature (T_Leaf_), net photosynthesis rate (A), carboxylation efficiency (CE), transpiration rate (E), stomatal conductance (g_sw_), and photosynthetic phosphorus and potassium use efficiencies (PPUE, PKUE). These traits show high similarity and are aggregated closely in the PCA biplot. A second cluster, shown in blue, comprises leaf water content (LWC), Rubisco nitrogen fraction (RNF), intercellular CO2 (Ci), and Rubisco use efficiency (RBUE), which appear more dispersed in the PCA but share underlying relationships. The third cluster, represented in green, includes leaf mass area (LMA), instantaneous and intrinsic water use efficiencies (WUE_ins_, WUE_int_), and photosynthetic nitrogen use efficiency (PNUE), which are positioned distinctly in the PCA biplot, particularly PNUE. This clustering pattern provides a structured view of the complex interactions among leaf traits, gas exchange parameters, and resource use efficiencies in *A. rugosa* under different light and nutrient levels, complementing the dimensional reduction achieved through PCA.

**Fig. 8.**
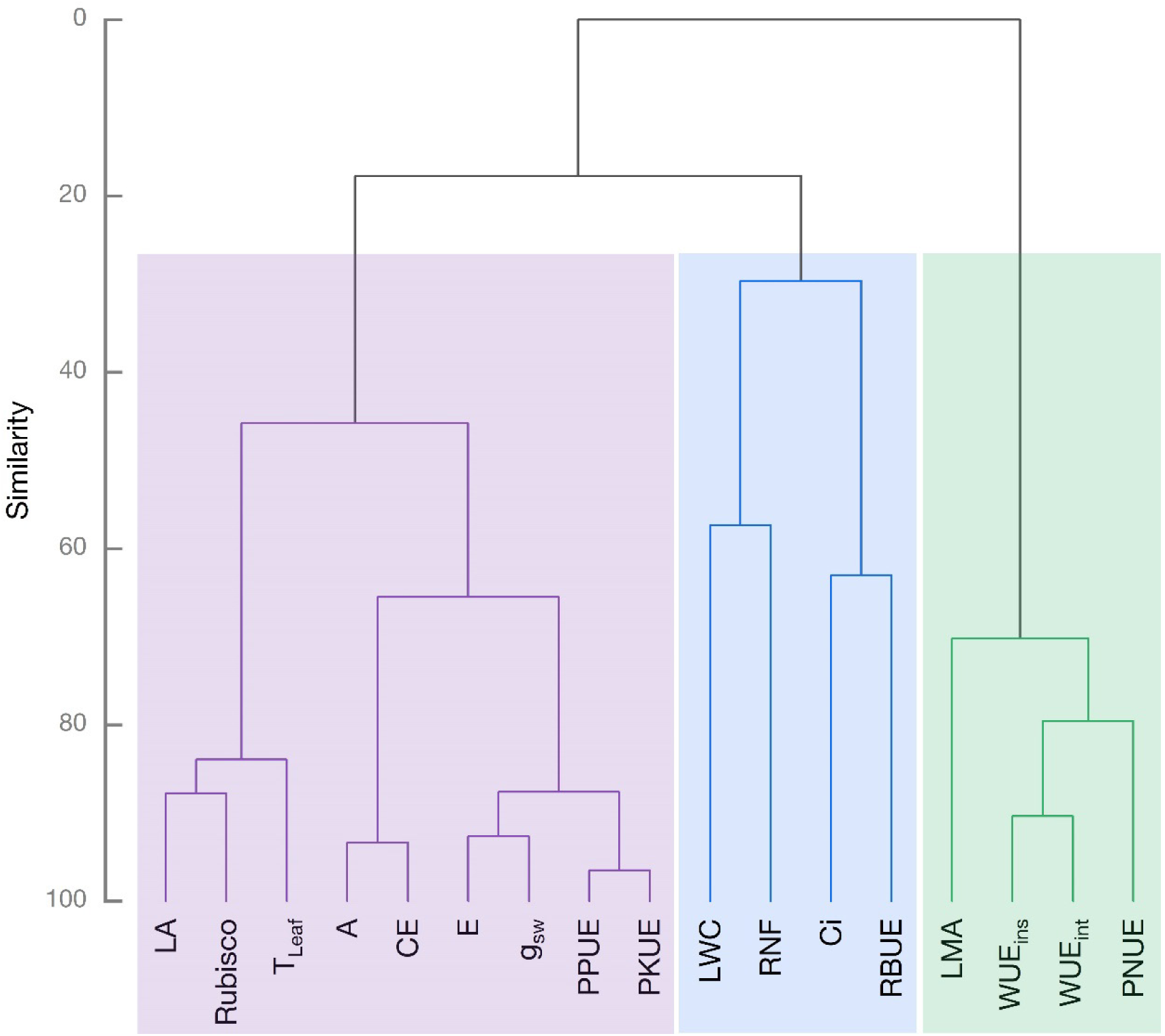
Clustering patterns among traits related to physiological plasticity and resource use in *A. rugosa* under different light and nutrient levels. Abbreviations: LA, leaf area; LMA, leaf mass area; LWC, leaf water content; A, net photosynthesis rate; E, transpiration rate; Ci, intercellular CO_2_; g_sw_, stomatal conductance; T_Leaf_, leaf temperature; CE, carboxylation efficiency; WUE_ins_, instantaneous water use efficiency; WUE_int_, intrinsic water use efficiency; RBUE, Rubisco use efficiency; RNF, Rubisco nitrogen fraction; PNUE, photosynthetic nitrogen use efficiency; PPUE, photosynthetic phosphorus efficiency; PKUE, photosynthetic potassium use efficiency. The traits in the boxes indicate that they have high similarity and are aggregated. The three boxes represent the four groups of clustering analysis results. Different colors are only used to better distinguish the three groups.

## Discussion

Our study on *A. rugosa* unveils a suite of adaptive responses to different light and nutrient levels, displaying remarkable physiological plasticity. Light availability was a dominant factor affecting the major leaf traits in *A. rugosa*. Plants grown under low-light conditions consistently exhibited larger leaf areas across all nutrient treatments, aligning with the well-established shade avoidance syndrome (Casal, 2012; Roig-Villanova and Martínez-García, 2016). This adaptation likely serves to maximize light interception in these low-light environments. However, our results also revealed a notable trade-off, as high-light conditions promoted greater leaf mass area (LMA). This pattern is consistent with the leaf economics spectrum theory, which posits that plants invest more in leaf structure and longevity under high-resource environments (Wright *et al*., 2004; Wang *et al*., 2023). The negative correlation between leaf area and LMA (r = -0.56, P < 0.001) further underscores this trade-off. Interestingly, our findings on leaf water content (LWC) deviate from some previous studies. We found higher LWC under low-light conditions, particularly pronounced at the highest nutrient level (NPK4). This contrasts with reports by Grunwald et al. (2024), who observed lower LWC in shaded inner-canopy across a broad range of species. Our results suggest that *A. rugosa* may employ a unique water retention strategy under low light, possibly to maintain cellular turgor and facilitate leaf expansion. Leaf temperature (TLeaf) showed minimal variation across treatments, with values ranging between 34°C and 37°C. However, a slight increase in T_Leaf_ was observed at higher nutrient levels, especially under low-light conditions. This relatively stable leaf temperature across treatments, despite variations in light exposure and nutrient levels, suggests that *A. rugosa* may possess some degree of thermoregulation. The capacity to sustain leaf temperatures within a narrow range, particularly under different light conditions, indicates subtle adaptive mechanisms that balance energy absorption, dissipation, and utilization (Michaletz *et al*., 2015, 2016). The positive correlation between leaf area and T_Leaf_ (r = 0.72, P < 0.001) suggests that larger leaves may face greater challenges in heat dissipation, potentially explaining the slight temperature increase in nutrient-rich, low-light conditions where leaf expansion was most distinct. This finding aligns with leaf energy balance, where larger leaves tend to have thicker boundary layers that can impede convective cooling (Leigh *et al*., 2017). The PCA results further elucidated these complex relationships, with leaf traits forming distinct clusters associated with light treatments. High-light treatments were strongly associated with increased LMA and water use efficiency, while low-light treatments correlated with larger leaf areas, higher photosynthetic rates, and slightly elevated leaf temperatures.

The interplay between light and nutrient levels profoundly affected the photosynthetic machinery and carbon assimilation in *A. rugosa*. Net photosynthesis rates significantly increased under high nutrient levels, particularly in HL-NPK4 treatment. This result supports the resource optimization theory predicting that plants allocate resources to maximize carbon acquisition when resources are abundant (Deans *et al*., 2020; Hartmann *et al*., 2020). However, we found an unexpected pattern in Rubisco dynamics. While Rubisco content generally increased with nutrient levels, low-light conditions increased Rubisco concentration across most nutrient treatments, except at the highest levels (NPK3 and NPK4). This contradicts the assumption that high-light environments stimulate greater investment in photosynthetic enzymes (Stitt and Schulze, 1994; Hikosaka *et al*., 2014). Our results suggest that *A. rugosa* may employ a compensatory mechanism in low light, increasing Rubisco content to maintain photosynthetic capacity despite reduced light energy. This is further supported by the patterns in Rubisco use efficiency (RBUE) and Rubisco nitrogen fraction (RNF), which were generally higher in low-light conditions. The strong positive correlation between net photosynthesis and both stomatal conductance (r = 0.82, P < 0.001) and carboxylation efficiency (r = 0.88, P < 0.001) suggests tight coordination between CO_2_ supply and demand. This may play a crucial role in the thermoregulatory capacity of *A. rugosa*, as stomatal control is a key mechanism for leaf temperature regulation (Chaves *et al*., 2016; Urban *et al*., 2017). The relationship between photosynthetic parameters and leaf temperature provides further insights into the species’ adaptive strategies. The slight increase in T_Leaf_ under high-nutrient, low-light environments coincided with elevated transpiration rates and stomatal conductance. This pattern indicates that *A. rugosa* may leverage increased nutrient levels to support greater transpirational cooling, particularly when light is limiting. Such a strategy can help sustain optimal leaf temperatures for photosynthetic processes, even in low-light environments where radiative heat load is decreased (Way and Yamori, 2014). The PCA revealed that RBUE aligned more closely with low-light treatments, suggesting that *A. rugosa* may optimize Rubisco utilization under shaded conditions. This optimization could be part of a strategy to maintain metabolic efficiency across a range of light environments, contributing to the species’ ecological plasticity.

Our analysis of resource use efficiencies provides novel insights into the adaptive strategies of *A. rugosa* under different light and nutrient levels. Water use efficiency (WUE) exhibited a complex response, with instantaneous WUE (WUE_ins_) being notably higher across all nutrient levels under high-light conditions. This finding is consistent with the general understanding that plants in high- light environments often display greater WUE to cope with higher evaporative demand (Syvertsen, 1984; Lambers *et al*., 2019; Ye *et al*., 2020). However, the intrinsic WUE (WUE_int_) response was more nuanced, displaying significant light-dependent differences only under low nutrient levels. This suggests that nutrient availability may modulate the plant’s water use strategy, particularly in resource-limited environments. The negative correlations between WUE indexes and most other physiological parameters, especially transpiration rate and stomatal conductance, underscore the trade-offs involved in optimizing water use versus carbon gain. These trade-offs likely play a vital role in the species’ thermoregulation, as maintaining high WUE can limit transpirational cooling (Carvalho *et al*., 2015). *A. rugosa* maintained relatively stable leaf temperatures despite changes in WUE, suggesting a balance between water storage and temperature regulation. Photosynthetic nutrient use efficiencies showed intriguing patterns across treatments. Photosynthetic nitrogen use efficiency (PNUE) was significantly higher in high-light conditions, consistent with the findings of Hikosaka and Terashima (1995), who reported increased PNUE in sun leaves. However, both photosynthetic phosphorus use efficiency (PPUE) and photosynthetic potassium use efficiency (PKUE) values were higher in low-light conditions, particularly at higher nutrient treatments. This unexpected result suggests that *A. rugosa* may have evolved mechanisms to optimize P and K use under low-light environments, possibly as an adaptation to understory growth. The robust positive correlation between PPUE and PKUE (r = 0.94, P < 0.001) shows coordinated regulation of these macronutrients in photosynthetic processes. The clustering of PPUE and PKUE with transpiration rate and stomatal conductance in both the correlation heatmap and PCA biplot suggests a tight coupling between nutrient use efficiency and leaf gas exchange properties. This coupling might also contribute to thermoregulation, as efficient nutrient utilization can support the maintenance of optimal photosynthesis and stomatal responses in various environmental conditions (Kumar *et al*., 2002; Sardans and Peñuelas, 2015). This capacity to maintain high PPUE and PKUE, in low-light conditions may represent a key adaptation to heterogeneous light environments, which allows the *A. rugosa* to optimize resource allocation and maintain physiological performance in shaded habitats.

The multivariate analyses conducted in this study provide a holistic view of the interactions among leaf traits, gas exchange parameters, and resource use efficiencies in *A. rugosa*. The PCA revealed distinct clustering patterns associated with light and nutrient treatments, explaining 62.9% of the total variation in the first two principal components. This high explanatory power highlights the strong effects of these environmental factors on the physio-biochemical responses of *A. rugosa*. The HCA further refined our understanding of trait relationships, identifying three major clusters of variables. The largest cluster, which consists of diverse parameters, including leaf area, Rubisco content, and various gas exchange parameters, underscores the integrated nature of plant responses to environmental changes. The inclusion of leaf temperature in this cluster, along with its positive correlation with leaf area and transpiration rate, supports the idea of coordinated thermoregulatory mechanisms involving multiple physiological and morphological traits (Michaletz *et al*., 2015). The clustering of WUE indexes with LMA and PNUE in a separate group emphasizes the distinct regulatory mechanisms governing water and nitrogen economics in *A. rugosa*. This division infers that the species may employ different strategies for optimizing water and nitrogen use, which can impact its ability to maintain physiological homeostasis across diverse environmental conditions. The third cluster, containing LWC, RNF, and intercellular CO2 concentration, points to a potential network between cellular water status, nitrogen allocation to photosynthetic machinery, and CO_2_ availability at the chloroplast level. This relationship may have also affected the species’ ability to maintain stable leaf temperatures and photosynthetic performance across treatments. Furthermore, these multivariate patterns provide a framework for understanding the coordinated plasticity of multiple traits in response to resource availability, supporting the concept of integrated phenotypes in plant adaptation (Pigliucci, 2003; Freschet *et al*., 2018). Our findings align with the growing recognition of phenotypic integration in mediating plant responses to environmental heterogeneity (Valladares *et al*., 2007; Matesanz *et al*., 2021). The observed correlations and clustering patterns suggest that *A. rugosa* employs a set of coordinated physiological and morphological adjustments to optimize resource acquisition and utilization under different light and nutrient conditions. This integrative approach to resource management, including the maintenance of thermal homeostasis, may contribute to the ecological success of *A. rugosa* and its potential for cultivation under diverse environmental conditions.

## Conclusions

In conclusion, our comprehensive study on *A. rugosa* reveals a suite of adaptive responses to light and nutrient variability, demonstrating the species’ high physiological plasticity. The light-nutrient interaction strongly affected leaf traits, photosynthetic machinery, and resource use efficiencies. Notably, *A. rugosa* showed unexpected patterns in Rubisco dynamics and nutrient use efficiencies, particularly in low light, suggesting a compensatory mechanism. The plant maintained relatively constant leaf temperatures across treatments, indicating subtle thermoregulatory adaptations that balance energy absorption, dissipation, and utilization. Our multivariate analyses unveiled distinct clustering patterns of physiological and morphological traits, highlighting the integrated nature of plant responses to environmental changes. These findings support our hypothesis of coordinated plasticity in leaf traits, photosynthetic processes, and resource-use efficiencies and provide new insights into the adaptive strategies of *A. rugosa* (Fig. 9.). The plant’s ability to optimize resource allocation and maintain physiological performance across light and nutrient gradients underscores its potential for cultivation under diverse conditions, particularly in the tropics, and contributes to our broader understanding of plant functional ecology in response to changing environments.

**Fig. 9.**
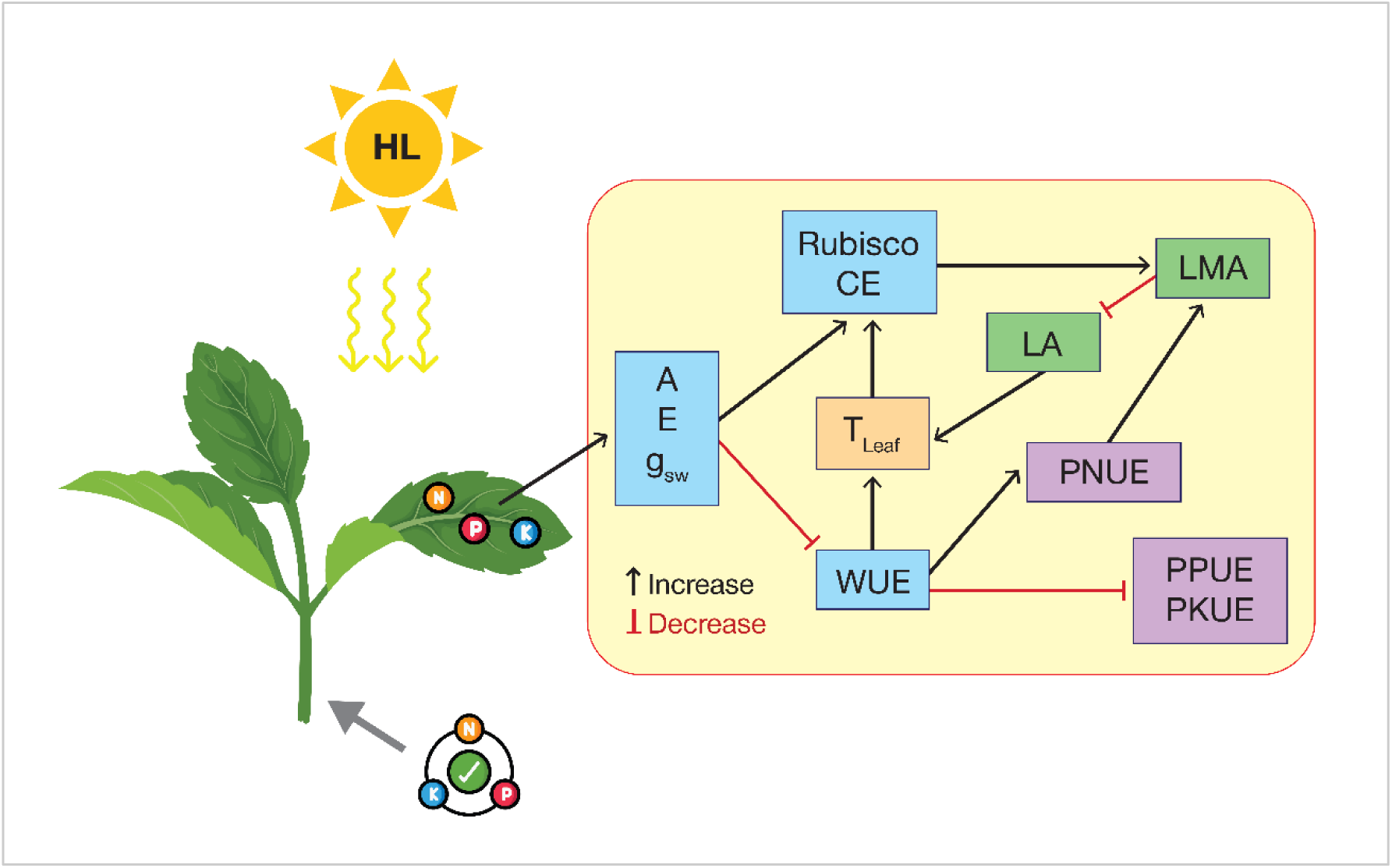
Schematic diagram illustrating the morphological and physiological adjustments in *A. rugosa* under high light and nutrient levels. Abbreviations: HL; high light, LA, leaf area; LMA, leaf mass area; A, net photosynthesis rate; E, transpiration rate; g_sw_, stomatal conductance; T_Leaf_, leaf temperature; CE, carboxylation efficiency; WUE, water use efficiency; WUE_int_, PNUE, photosynthetic nitrogen use efficiency; PPUE, photosynthetic phosphorus efficiency; PKUE, photosynthetic potassium use efficiency.

## Author Contributions

**Khairul Azree Rosli:** Conceptualization, Methodology, Experimentalize, Data analysis, Writing- original draft preparation. **Puteri Edaroyati Megat Wahab:** Conceptualization, Methodology, Revision, Supervision. **Azizah Misran:** Revision, Supervision. **Latifah Saiful Yazan:** Revision, Supervision. All authors have read and revised the manuscript, provided helpful discussions, and approved its final version.

## Data availability

The data and materials supporting this study’s findings are available from the corresponding author upon request.

## Acknowledgements

This work has received funding from the Putra Graduate Initiative Grant (GP-IPS) from Universiti Putra Malaysia (Vot 9660900). We also acknowledge other researchers and staff for their excellent management and general skills.

